# Cell growth and nutrient availability control the mitotic exit signaling network in budding yeast

**DOI:** 10.1101/2023.08.04.552008

**Authors:** Rafael A. Talavera, Beth E. Prichard, Robert A. Sommer, Ricardo M. Leitao, Christopher J. Sarabia, Semin Hazir, Joao A. Paulo, Steven P. Gygi, Douglas R. Kellogg

## Abstract

Cell growth is required for cell cycle progression. The amount of growth required for cell cycle progression is reduced in poor nutrients, which leads to a reduction in cell size. In budding yeast, nutrients influence cell size by modulating the duration and extent of bud growth, which occurs predominantly in mitosis. However, the mechanisms are unknown. Here, we used mass spectrometry to identify proteins that mediate the effects of nutrients on bud growth. This led to the discovery that nutrients regulate numerous components of the Mitotic Exit Network (MEN), which controls exit from mitosis. A key component of the MEN undergoes gradual multi-site phosphorylation during bud growth that is dependent upon growth and correlated with the extent of growth. Furthermore, activation of the MEN is sufficient to over-ride a growth requirement for mitotic exit. The data suggest a model in which the MEN integrates signals regarding cell growth and nutrient availability to ensure that mitotic exit occurs only when sufficient growth has occurred.

## Introduction

In all orders of life, cell cycle progression is dependent upon cell growth, which ensures that dividing cells attain sufficient volume and mass so that cell division produces viable cells of an appropriate size (Jorgensen and Tyers, 2004; Turner *et al*., 2012; Ginzberg *et al*., 2015). The threshold amount of growth required for cell cycle progression is reduced in poor nutrients, which leads to a large reduction in cell size (Kellogg and Levin, 2022). The reduced growth threshold in poor nutrients likely confers a competitive advantage, as it allows cells to divide and proliferate more rapidly when nutrients become limiting. The mechanisms that set and modulate the threshold amount of growth required for cell cycle progression are largely unknown.

In budding yeast, most cell growth takes place during growth of the daughter bud, which occurs almost entirely during mitosis (Leitao and Kellogg, 2017). Bud growth is required for progression through mitosis, and nutrient availability strongly influences the rate, duration and extent of bud growth (Anastasia *et al*., 2012; Leitao and Kellogg, 2017). Thus, in poor nutrients the rate of bud growth is reduced, yet the duration of bud growth is extended, presumably to allow more time for bud growth. In effect, the threshold amount of bud growth required for completion of mitosis is reduced, leading to birth of very small daughter cells. These observations suggest the existence of nutrient and growth-sensing mechanisms that modulate the duration and extent of bud growth to ensure that daughter cells are born at an appropriate size.

As the bud grows during mitosis, approximately equal volumes are added in metaphase and anaphase (Leitao and Kellogg, 2017). Nutrient availability modulates the duration and extent of bud growth in both intervals, and recent work suggests that the two intervals are controlled by different mechanisms (Leitao *et al*., 2019; Jasani *et al*., 2020). The metaphase interval is influenced by a pair of related kinases called Gin4 and Hsl1. Both kinases are required for normal control of bud growth and undergo gradual hyperphosphorylation and activation that appear to be dependent upon bud growth and proportional to the extent bud growth, which suggests that they play roles in measuring growth (Jasani *et al*., 2020)5/2/23 2:57:00 PM. Once they have been fully activated, Gin4 and Hsl1 promote mitotic progression by inhibiting Swe1, the budding yeast homolog of the conserved Wee1 kinase that directly phosphorylates and inhibits mitotic Cdk1/cyclin complexes (Ma *et al*., 1996; Longtine *et al*., 2000). In both mammalian cells and budding yeast, it is thought that Wee1 keeps mitotic Cdk1 activity low during metaphase and that inhibition of Wee1 leads to a full activation of Cdk1 that drives the metaphase to anaphase transition (Deibler and Kirschner, 2010; Harvey *et al*., 2011). Gin4 and Hsl1 are required for normal control of the duration and extent of bud growth in metaphase but are not required during anaphase/telophase (Leitao *et al*., 2019; Jasani *et al*., 2020). These data suggest that growth in anaphase/telophase is regulated by mechanisms that are distinct from those that work in metaphase. The mechanisms by which nutrients modulate the duration and extent of bud growth in anaphase are unknown.

Here, we used proteome-wide mass spectrometry to search for proteins that influence the duration and extent of bud growth in late mitosis. The results point to the Mitotic Exit Network (MEN), an essential signaling network that includes highly conserved proteins, as a likely target of signals that link mitotic exit to cell growth and nutrient availability. Previous studies suggested that the MEN links mitotic exit to mitotic spindle orientation and elongation (Bardin *et al*., 2000; Pereira *et al*., 2000; Campbell *et al*., 2020). Our studies establish for the first time that key components of the MEN also respond to signals associated with bud growth and nutrient availability.

## Results

### Multiple components of the mitotic exit network are controlled by nutrient-dependent signals

Previous studies found that nutrients modulate the duration and extent of bud growth in both metaphase and anaphase, and that changes in nutrient availability immediately influence cell cycle progression and the threshold amount of growth required for cell cycle progression (Fantes and Nurse, 1977; Leitao and Kellogg, 2017). The data suggest that a reduced growth rate caused by a shift to poor nutrients triggers an immediate mitotic delay to allow more time for bud growth (Leitao and Kellogg, 2017). Therefore, to identify signals that could play a role in modulating the duration and extent of growth in mitosis, we used proteome-wide mass spectrometry to search for proteins that undergo large changes in phosphorylation in response to a rapid shift from rich to poor carbon during mitosis. We reasoned that these could include proteins that play roles in setting the threshold amount of bud growth required for cell cycle progression.

Cells growing in medium containing a rich carbon source (YP + 2% dextrose) were released from a G1 arrest and rapidly shifted to poor carbon medium (YP + 2% glycerol/2% ethanol) at 90 minutes, when the mitotic cyclin Clb2 reached peak levels (**Figure 1**). Proteins were isolated from the shifted and unshifted control cells 10 minutes after the shift and proteolytic peptides were analyzed by quantitative proteome-wide mass spectrometry to search for changes in phosphorylation. The ratio of phosphorylated to unphosphorylated peptides in shifted versus unshifted control cells was log2 transformed. Thus, a negative log2 ratio indicates a loss of phosphorylation in response to a shift to poor carbon, whereas a positive log2 ratio indicates a gain of phosphorylation. A total of three biological replicates were analyzed, which allowed calculation of average log2 ratios and standard deviation. The complete data set is shown in Table S1.

**Figure 1.**
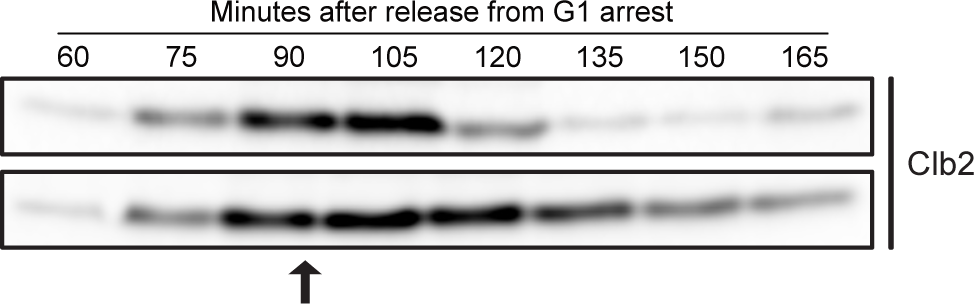
– Analysis of the effects of a shift from rich to poor carbon during metaphase. Cells were grown overnight in YPD medium and arrested in G1 phase with alpha factor. The cells were then released from the arrest into fresh YPD at 25°C. At the point indicated by the arrow, half of the culture was shifted to poor carbon (bottom panel). Samples were collected at the indicated time points to analyze Clb2 levels by western blot. Samples for mass spectrometry analysis were collected ten minutes after the shift to poor carbon.

We searched the mass spectrometry data for proteins involved in mitotic progression and cell growth, which are summarized in Table 1. A log2 ratio of +/- 1.32, corresponding to a 2.5-fold change in phosphorylation occupancy, was used as an arbitrary cutoff to define substantial changes in phosphorylation. Numerous components of a signaling network that regulates exit from mitosis, referred to as the “mitotic exit network” (MEN), were amongst the proteins that showed the largest loss of phosphorylation when cells were shifted from rich to poor nutrients in mitosis. These proteins include Net1, Cdc14, Lte1, Cdc15, Bfa1, Nud1, and Zds1. Previous work suggested that the MEN plays a central role in triggering mitotic exit by initiating destruction of the mitotic cyclin Clb2 and inhibition of mitotic Cdk activity (Visintin *et al*., 1998, 1999; Jaspersen *et al*., 1999; Shou *et al*., 1999; Stegmeier *et al*., 2002). It has been proposed that the MEN initiates mitotic exit in response to signals generated by proper orientation of the mitotic spindle in the daughter bud (Bardin *et al*., 2000; Pereira *et al*., 2000). Here, the discovery that numerous components of the mitotic exit network show large changes in phosphorylation in response to a shift to poor nutrients indicates that the MEN is strongly regulated by nutrient-dependent signals. Since the MEN can influence the duration of late mitosis, the data further suggest that the MEN could be the target of nutrient-dependent signals that influence the duration and extent of bud growth in late mitosis.

**Table 1.**
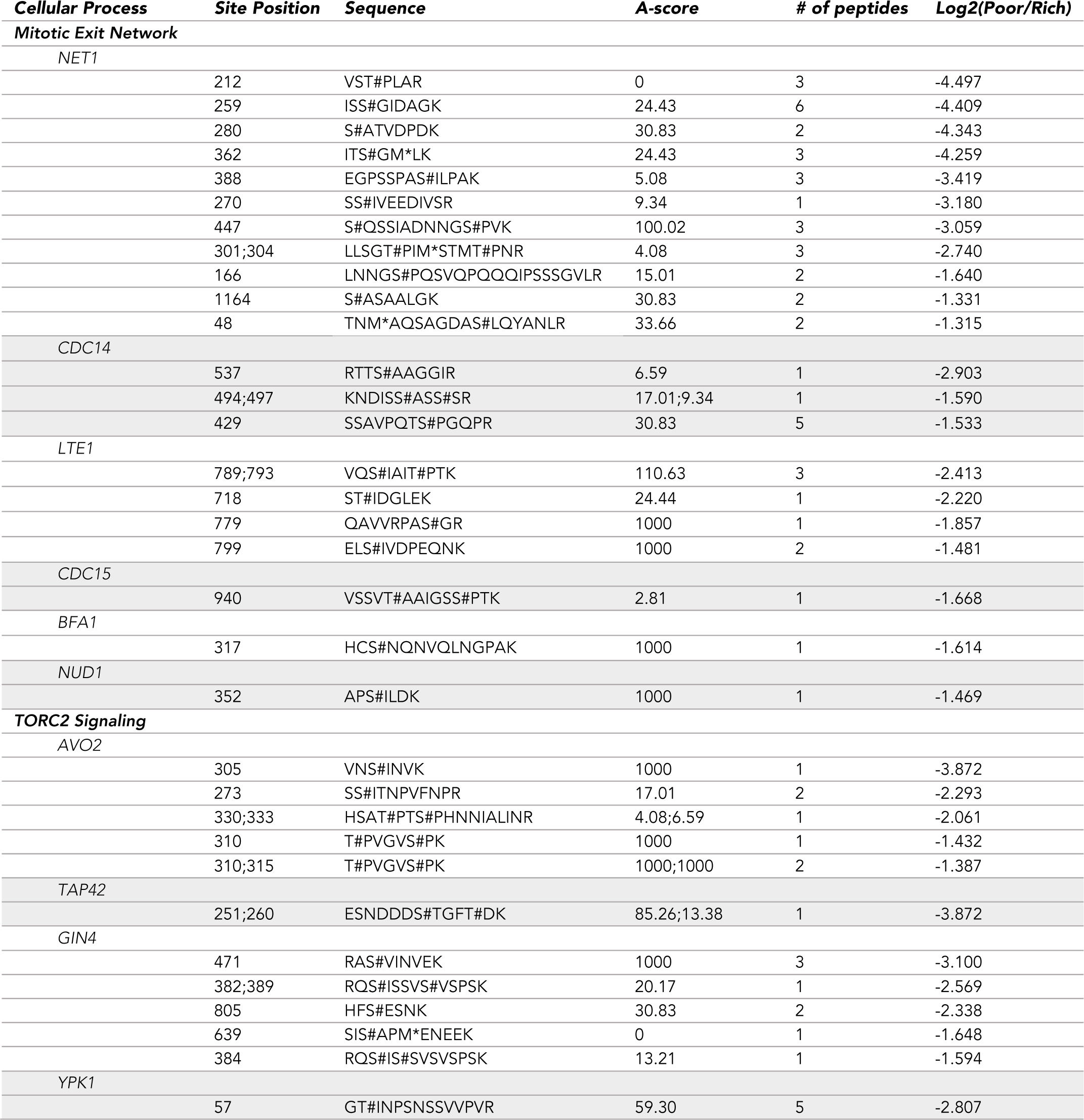
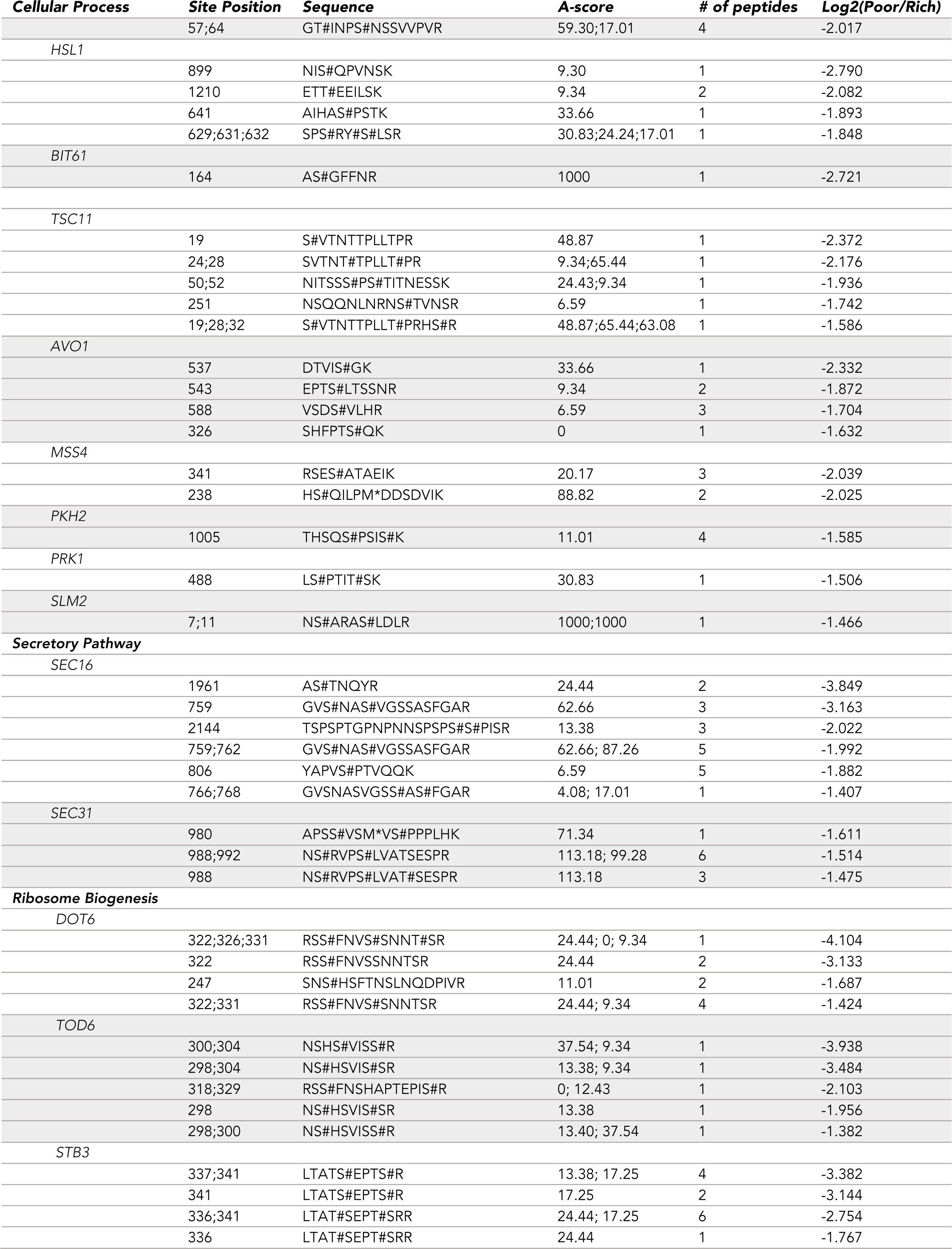
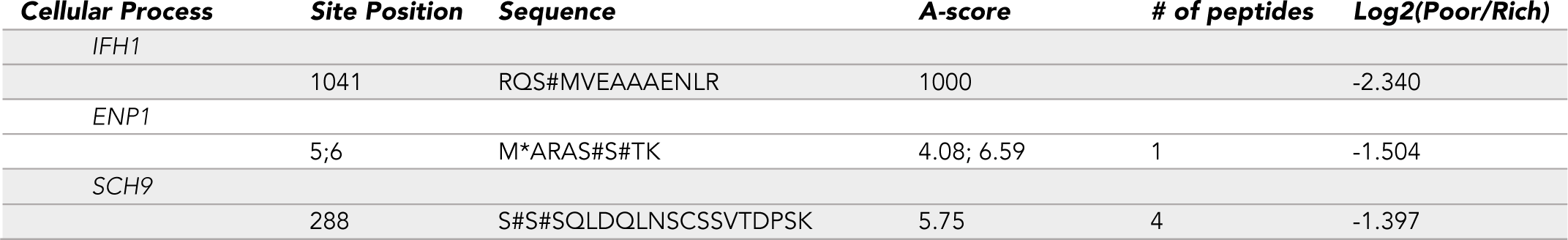
– A summary of key peptides that undergo substantial changes in phosphorylation in response to a shift to poor carbon during mitosis. The A-score indicates the likelihood a phosphorylation site is correctly localized within a peptide. A given phosphorylation site is correctly assigned with >95% confidence when the A-score is >13. An A-score of 0 indicates that the phosphorylated residue within the peptide sequence cannot be precisely determined. An asterisk (*) represents oxidation of the preceding methionine, an artifact of the sample preparation. A hashtag indicates that the preceding amino acid is likely the relevant phosphorylation site.

The mass spectrometry analysis also identified multiple components of TOR Complex 2 (TORC2) and its surrounding signaling network (Table 1). These proteins include Avo2, Tsc11, Ypk1, Mss4, Pkh2, Gin4, Hsl1, Rts1, and Prk1. The identification of components of the TORC2 network is consistent with previous studies that found that TORC2 signaling is required for nutrient modulation of cell size and growth rate (Lucena *et al*., 2018). As an example, loss of Rts1, a regulatory subunit for PP2A, causes misregulation of TORC2 signaling, as well as a complete loss of the proportional relationship between cell size and growth rate during bud growth (Lucena *et al*., 2018; Leitao *et al*., 2019).

Finally, the analysis identified several core components of the COPII complex that is required for ER to Golgi transport (Sec16 and Sec31) as well as a key regulator of ribosome biogenesis (Sch9) (Table 1). Since a shift to poor nutrients leads to a rapid reduction in growth rate, these findings suggest that poor nutrients trigger signals that rapidly reduce the rates of both ribosome biogenesis and membrane growth to match reduced biosynthetic rates in poor nutrients.

### Net1 hyperphosphorylation occurs before mitotic exit

The discovery that multiple components of the MEN are regulated in response to changes in carbon source suggested that the MEN could play a role in linking mitotic exit to cell growth and nutrient availability. In this context, the discovery that Net1 phosphorylation is strongly modulated by carbon source was particularly interesting. Net1 showed large decreases in phosphorylation at 11 sites, including one site that has not been detected in any previous studies. Net1 was also identified as a top hit in a mass spectrometry screen for proteins that undergo large changes in phosphorylation in response to an arrest of plasma membrane growth during early mitosis (Clarke *et al*., 2017). Net1 plays a critical role in initiation of mitotic exit and also controls ribosome biogenesis, which is a major facet of cell growth (Shou *et al*., 1999, 2001; Straight *et al*., 1999; Visintin *et al*., 1999; Shou and Deshaies, 2002; Hannig *et al*., 2019). Net1 is localized in the nucleolus early in the cell cycle, where it binds and inhibits Cdc14 (Shou *et al*., 1999; Visintin *et al*., 1999; Traverso *et al*., 2001). Hyperphosphorylation of Net1 is thought to be a critical step that helps release Cdc14 from Net1 so that it can leave the nucleolus to initiate mitotic exit (Shou *et al*., 1999; Yoshida and Toh-e, 2002; Visintin *et al*., 2003). In previous work we found evidence that the events of bud growth generate signals that are dependent upon growth and proportional to growth (Anastasia *et al*., 2012; Jasani *et al*., 2020). We therefore considered a working hypothesis in which the events of bud growth generate a signal that contributes to Net1 phosphorylation and helps trigger activation of the MEN, while nutrient-dependent signals modulate the strength of the growth-dependent signal needed to initiate mitotic exit.

Since hyperphosphorylation of Net1 is thought to be a key early step in the initiation of mitotic exit, we carried out new experiments to determine whether Net1 phosphorylation is influenced by nutrients or growth, which would begin to test the hypothesis that the MEN links mitotic exit to cell growth. Previous studies reported that hyperphosphorylation of Net1 can be detected via electrophoretic mobility shifts; however, we found that the large size of Net1 made reproducible detection of these shifts difficult. We therefore developed an additional tool for detection of Net1 phosphorylation. We reasoned that expression of a truncated version of Net1 could provide a readout of phosphorylation while also allowing easier detection due to the reduced size of the truncated Net1. We analyzed proteome-wide mass spectrometry data from multiple studies that have been collated on BioGrid, which indicated that most phosphorylation of Net1 occurs in the first two thirds of the protein. We therefore constructed a plasmid that expresses a version of Net1 that lacks the last 328 amino acids and is tagged with 3xHA (Net1^Δ328^-3xHA). The truncated Net1 is expressed from the endogenous promoter and can be integrated at URA3 so that cells express both full-length Net1 as well as the truncated reporter. The reporter includes an N-terminal Cdc14 binding site that is thought to undergo phosphorylation that helps release Cdc14 (Traverso *et al*., 2001).

Current models suggest that phosphorylation of Net1 is a critical event that helps initiate mitotic exit, which would suggest that Net1 phosphorylation is initiated late in mitosis during mitotic exit. However, in a previous study it appears that phosphorylation of Net1 is initiated early in mitosis as mitotic cyclin levels are rising, which could suggest that Net1 phosphorylation is initiated before mitotic exit (Visintin *et al*., 2003). To further investigate, we tested whether arresting cells at metaphase, before mitotic exit, blocked hyperphosphorylation of Net1. To do this, we released cells from a G1 phase arrest into YPD medium containing benomyl, which depolymerizes microtubules and activates a mitotic spindle checkpoint that arrests cells before metaphase. Depolymerization of microtubules will also eliminate any signals generated by proper orientation of the mitotic spindle, which is thought to be a critical trigger for mitotic exit. As a control, we also released cells from the arrest into YPD medium that does not contain benomyl. Phosphorylation of both full length Net1 and Net1^Δ328^ was analyzed by western blot to detect electrophoretic mobility shifts, and levels of the mitotic cyclin Clb2 were analyzed as a molecular marker for mitotic progression. In the control cells without benomyl, both Net1 and Net1^Δ328^ underwent hyperphosphorylation as cells progressed through mitosis, with peak phosphorylation appearing to occur shortly after peak Clb2 levels, which likely corresponds to anaphase/telophase (**Figure 2A,B**). Both Net1 and Net1^Δ328^ underwent extensive hyperphosphorylation when cells were arrested before mitotic exit with benomyl. The extent of Net1 hyperphosphorylation was greater in benomyl-arrested cells compared to the control cells, which will be addressed further below.

**Figure 2.**
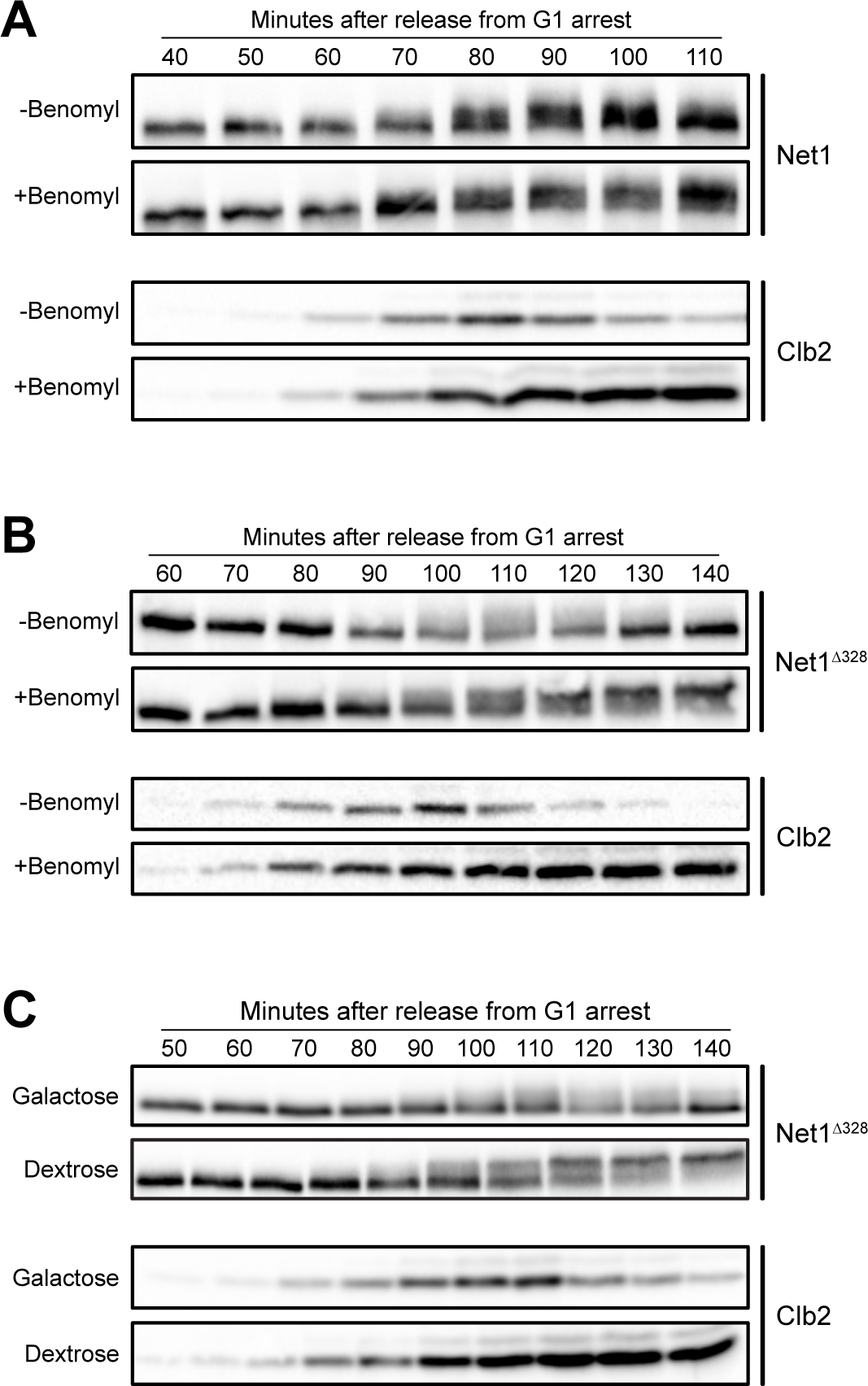
– Net1 hyperphosphorylation occurs before mitotic exit. *NET1-6xHA* (A,B) or *NET1^Δ328^-3xHA* (C,D) cells growing in YPD were released from a G1 arrest at 25°C into YPD or YPD containing benomyl. Samples were collected at the indicated time points to assay Net1 phosphorylation and Clb2 protein levels by western blot. Alpha factor was added back to the cultures at 60 minutes after release to prevent a second cell cycle. (C) *NET1^Δ328^-3xHA GAL1-CDC20* cells grown overnight to log phase in YPGal at room temperature were released from a G1 phase arrest into YPGal or YPD at 25°C. Samples were collected at the indicated time intervals to assay for net1^Δ 328^ and Clb2 by western blot.

To further test whether Net1 phosphorylation occurs before mitotic exit we used depletion of Cdc20 as an alternative means of arresting cells in metaphase (**Figure 2C)**. Again, Net1 underwent full hyperphosphorylation in metaphase-arrested cells.

These results show that hyperphosphorylation of Net1 that can be detected via electrophoretic mobility shifts does not depend on the events of mitotic exit or any functions of the mitotic spindle.

### Net1 phosphorylation is influenced by carbon source

We next analyzed whether Net1 phosphorylation is influenced by carbon source. In previous studies we showed that the durations of both metaphase and anaphase are increased in cells growing in poor carbon, while the extent of bud growth during both intervals is reduced (Leitao and Kellogg, 2017). Cells growing in rich or poor carbon were released from a G1 arrest and Net1 phosphorylation was assayed by western blot (**Figure 3**). Clb2 levels were analyzed in the same samples to provide a marker for mitotic progression. In rich carbon, both full length Net1 and Net1^Δ328^ underwent hyperphosphorylation during mitosis. In poor carbon, the extent of Net1 phosphorylation was reduced and the duration of Net1 phosphorylation was extended. Since the rate and extent of bud growth are strongly reduced in poor carbon, these observations suggest that extent of Net1 phosphorylation is correlated with the rate and extent of bud growth. Previous work found that Gin4 and Hsl1, which are thought to relay growth-dependent signals before mitotic exit, also undergo hyperphosphorylation that is correlated with the rate and extent of bud growth in early in mitosis (Jasani *et al*., 2020).

**Figure 3.**
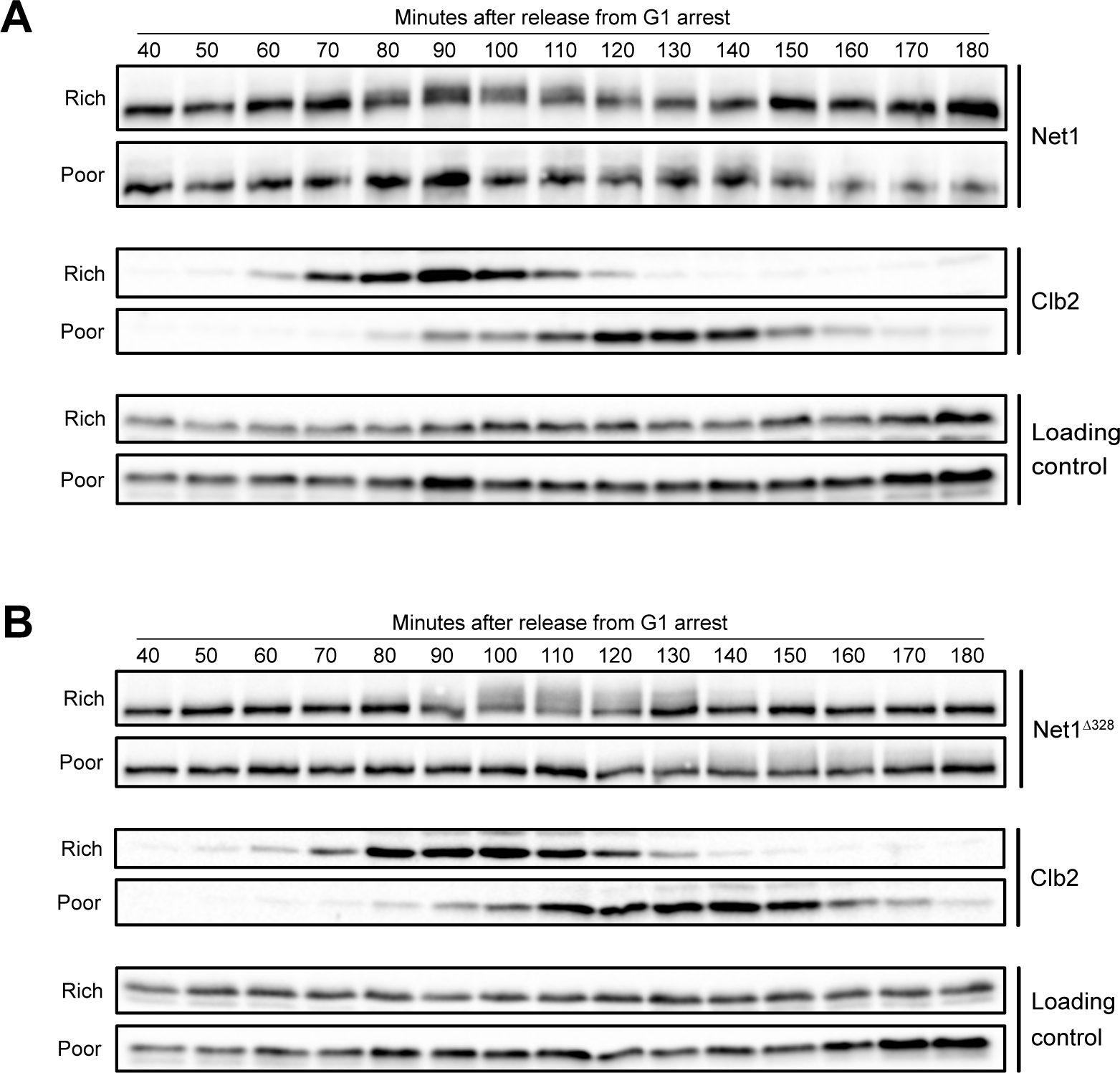
– Net1 hyperphosphorylation is reduced in poor carbon. *NET1-6xHA* (A) or *NET1^Δ328^-3xHA* (B) cells grown overnight in YPD or YPG/E were released from a G1 phase arrest into fresh medium at 25°C Samples were collected at the indicated time points to assay for Net1 and Clb2 by western blot. An anti-Nap1 antibody was used for a loading control. Alpha factor was added back to the cultures 60 minutes after release to prevent a second cell cycle.

### Hyperphosphorylation of Net1 is correlated with the rate and extent of bud growth

To further investigate the relationship between Net1 phosphorylation and cell growth, we took advantage of the fact that bud growth continues during a prolonged mitotic arrest, which leads to growth of unusually large buds as well as an unusually long interval of bud growth (Gihana *et al*., 2021). We also took advantage of the fact that the rate and extent of bud growth in mitosis can be modulated by the quality of the carbon source present in the culture medium (Leitao and Kellogg, 2017). Thus, we tested whether the extent of Net1 hyperphosphorylation was correlated with growth when bud growth was prolonged and increased by a benomyl arrest, and when the rate of growth during the arrest was modulated by carbon source.

Cells were grown in carbon sources of varying quality and released from a G1 phase arrest into media containing benomyl to arrest the cell cycle before metaphase. The cells were then maintained at the arrest point for a prolonged interval of time. We utilized dextrose as a rich carbon source, galactose as an intermediate quality carbon source, and glycerol/ethanol as a poor carbon source. Samples were taken at regular intervals during bud growth and phosphorylation of the Net1^Δ328^ reporter was measured by western blot (**Figure 4A**) and median cell size was measured with a Coulter Counter. Cell size was plotted as a function of time to provide a measure of the rate and extent of bud growth (**Figure 4B**).

**Figure 4.**
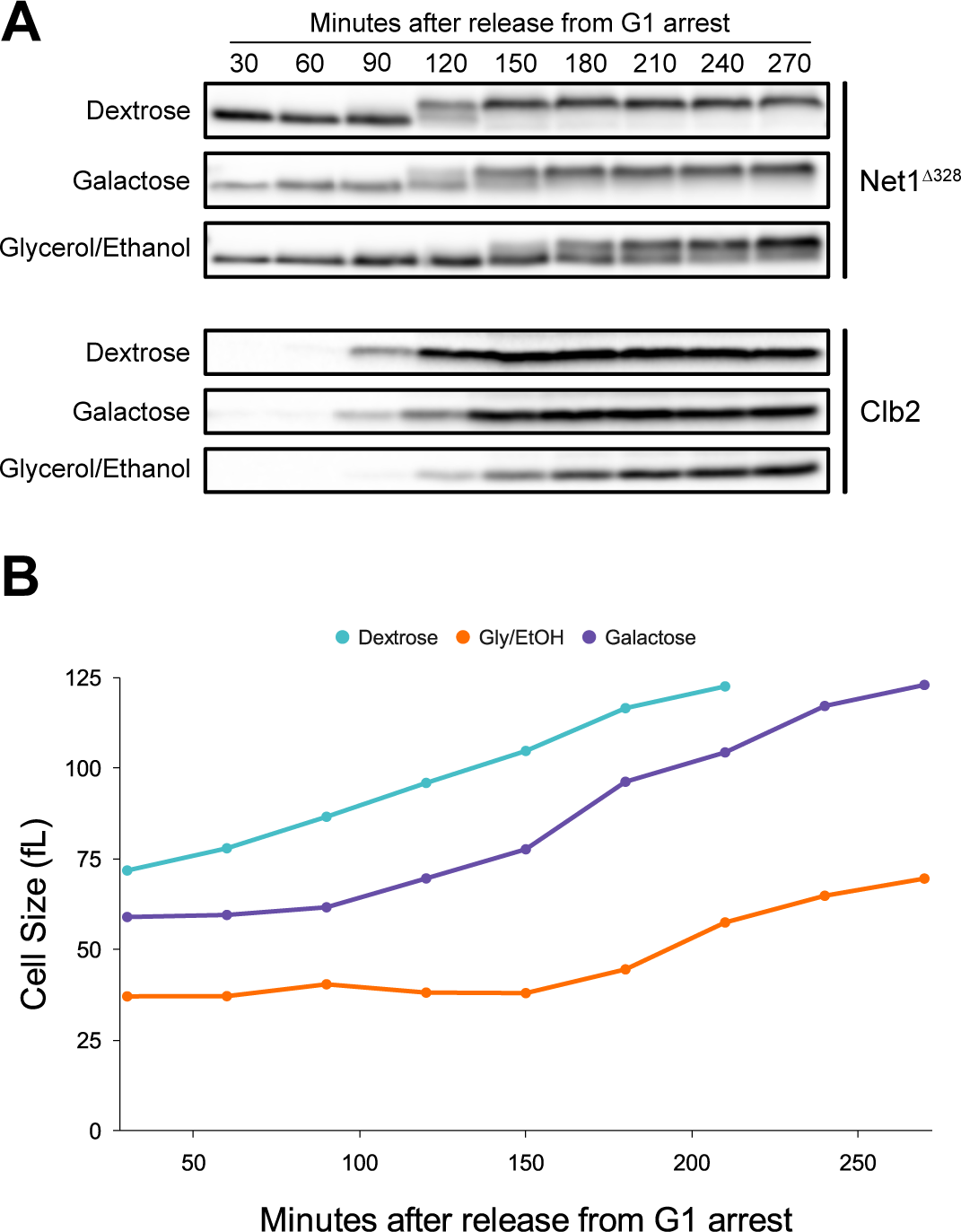
– Hyperphosphorylation of Net1 is correlated with the rate and extent of bud growth. (A) *NET1^Δ328^-3xHA* cells grown overnight to log phase in YPD, YPGal, or YPG/E were released from a G1 phase arrest at 25°C into fresh media containing benomyl. Samples were taken at the indicated times to assay for Net1^Δ328^-3xHA and Clb2. An anti-Nap1 antibody was used for a loading control. (B) Samples from the same cultures in panel A were fixed at the indicated time intervals and median cell size was measured using a Coulter Counter.

In rich carbon, buds underwent rapid and extensive growth. Net1^Δ328^ phosphorylation appeared to go to completion during the prolonged arrest, and the maximal extent of Net1^Δ328^ phosphorylation achieved during the prolonged benomyl arrest was greater than in cells going through a normal cell cycle (compare top panels in **Figures 4A** and **2B**). Since daughter buds grow to be much larger in benomyl-arrested cells, this observation suggests that the extent of Net1 phosphorylation is correlated with the extent of bud growth. In galactose, the rate of bud growth was reduced and it took longer for Net1^Δ328^ phosphorylation to go to completion. In glycerol/ethanol, the poorest carbon source, the rate and extent of bud growth were strongly reduced and Net1 failed to reach full hyperphosphorylation. Together, these results provide further evidence that Net1 phosphorylation is strongly correlated with the rate and extent of bud growth.

### Hyperphosphorylation of Net1 is dependent upon membrane trafficking events that drive bud growth

To further investigate the relationship between Net1 phosphorylation and bud growth, we tested whether hyperphosphorylation of Net1 is dependent upon bud growth. In previous work, we tested for dependency of signaling events on bud growth by blocking plasma membrane growth in the daughter bud, which can be achieved by inactivating Sec6, a component of the exocyst complex that is required for fusion of vesicles with the plasma membrane during bud growth (Anastasia *et al*., 2012). A temperature sensitive allele of *SEC6* (*sec6-4*) can be used to achieve rapid conditional inactivation of Sec6.

Inactivation of Sec6 before bud emergence causes cells to arrest at metaphase, due most likely to a failure in bud growth during early mitosis (Anastasia *et al*., 2012). Swe1, the budding yeast homolog of Wee1, is required for the arrest and is thought to respond to signals that measure bud growth before metaphase. Loss of Swe1 allows cells that lack Sec6 function to proceed through metaphase to late anaphase/telophase, and the cells show normal mitotic spindle assembly and chromosome segregation within the mother cell (Anastasia *et al*., 2012). Therefore, we analyzed Net1 phosphorylation in wild type and *sec6-4 swe1Δ* cells, which ensured that any effects on Net1 phosphorylation were not due to a metaphase arrest. Cells were released from a G1 phase arrest and shifted to the restrictive temperature at 20 minutes after release from the arrest, before bud emergence. Hyperphosphorylation of full length Net1 was assayed by western blot and levels of Clb2 were assayed as a marker for mitotic progression. Hyperphosphorylation of Net1 failed to occur when *sec6-4* was inactivated (**Figure 5**). Controls showed that *sec6-4* alone also caused a failure in Net1 phosphorylation, while *swe1Δ* alone had no effect on Net1 phosphorylation (not shown). Thus, hyperphosphorylation of Net1 is independent of mitotic exit and mitotic spindle function but is completely dependent upon membrane trafficking events that are required for bud growth. Furthermore, the mitotic cyclin Clb2 accumulates to high levels in *sec6-4 swe1Δ* cells and is capable of driving mitotic spindle assembly and segregation of chromosomes in anaphase (Anastasia *et al*., 2012), which indicates that Clb2/Cdk1 activity is not sufficient to drive Net1 phosphorylation. This finding is consistent with a previous study that found that Clb2/Cdk1 activity likely plays a minor role in regulation of Net1, but only after MEN signaling has been initiated (Azzam *et al*., 2004).

**Figure 5.**
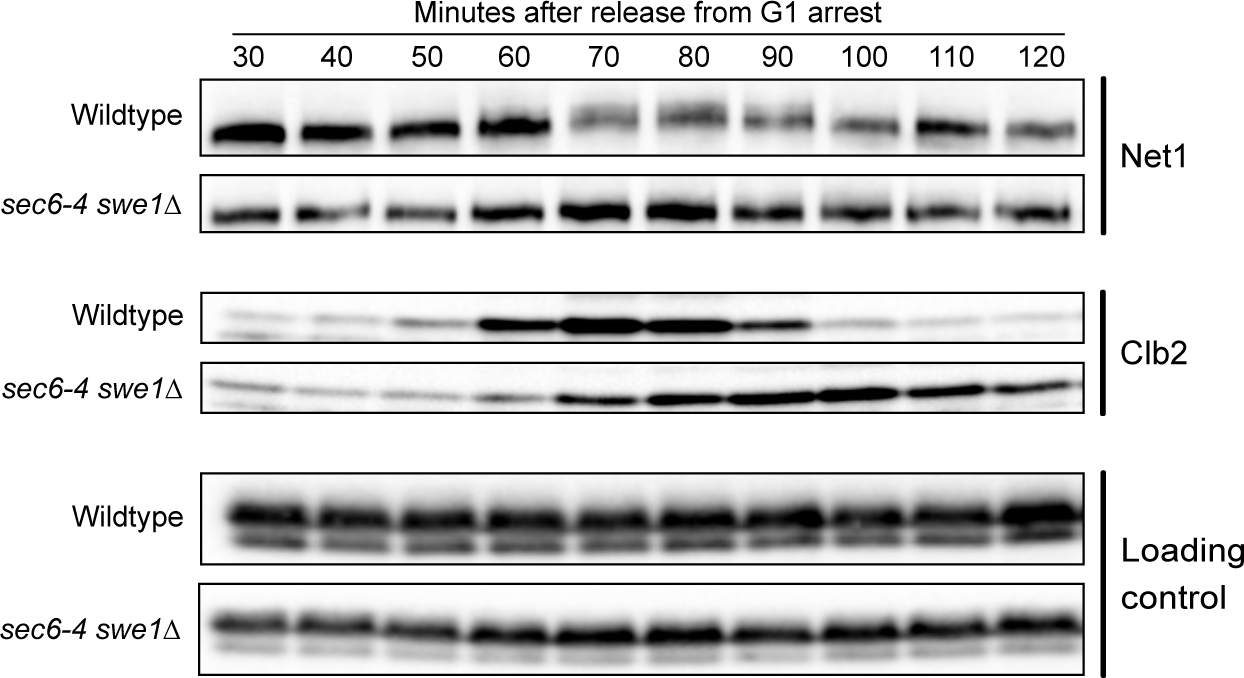
- Net1 hyperphosphorylation is dependent upon membrane trafficking events that are required for bud growth. *NET1^Δ328^-3xHA* and *NET1^Δ328^-3xHA sec6-4 swe1Δ* cells grown overnight to log phase in YPD were released from a G1 phase arrest at room temperature and were then shifted to the restrictive temperature (34°C) 20 minutes after release. Samples were taken at the indicated time intervals to assay for Net1^Δ328^-3xHA, Clb2 by western blot. An anti-Nap1 antibody was used for a loading control.

### An allele of *CDC14* that is not inhibited by Net1 bypasses the anaphase arrest caused by a failure in bud growth

The data thus far suggest that hyperphosphorylation of Net1 is dependent upon bud growth and correlated with the extent of bud growth. To explain the data, we hypothesized that hyperphosphorylation of Net1 is driven by growth-dependent signals. We further hypothesized that growth-dependent hyperphosphorylation of Net1 plays a role in activation of the MEN, which would help ensure that mitotic exit occurs only when sufficient growth has occurred. This model suggests that the late anaphase arrest in *sec6-4 swe1Δ* cells is caused by a failure in the release of Cdc14 from Net1. To test this model, we determined whether an allele of Cdc14 that shows reduced binding to Net1 (*cdc14^TAB6^*) is sufficient to bypass the late mitotic arrest in *sec6-4 swe1Δ* cells. The *cdc14^TAB6^* allele was identified in a screen for mutants that bypass a telophase arrest caused by loss of Cdc15 function, which indicates that the allele can bypass a failure to activate the MEN (Shou *et al*., 2001; Shou and Deshaies, 2002). We released *sec6-4 swe1Δ cdc14^TAB6^* cells and control cells from a G1 phase arrest at the permissive temperature and then shifted to the restrictive temperature 20 minutes later and assayed Clb2 protein levels by western blot to assess mitotic progression (**Figure 6A**). The *cdc14^TAB6^* allele fully abrogated the late anaphase arrest caused by *sec6-4*.

**Figure 6.**
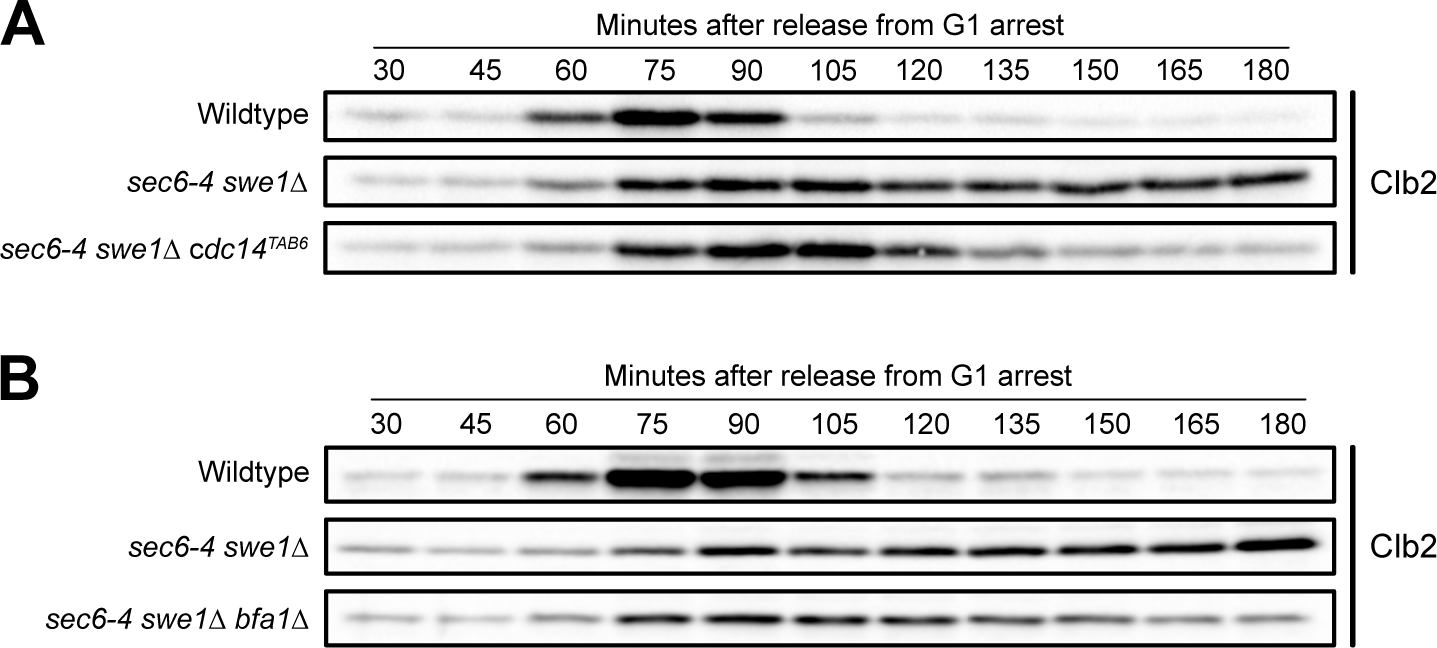
– An allele of *CDC14* that is not inhibited by Net1 bypasses the anaphase arrest caused by a failure in bud growth. (A) *NET1-6xHA*, *NET1-6xHA sec6-4 swe1Δ,* and *NET1-6xHA sec6-4 swe1Δ cdc14^TAB6^* cells grown overnight in YPD to log phase were released from a G1 phase arrest into fresh media at 25°C. 20 minutes after release the cultures were shifted to 34°C to inactivate sec6-4 and samples were collected at the indicated time points to assay levels of Clb2. (B) *NET1-6xHA*, *NET1-6xHA sec6-4 swe1Δ,* and *NET1-6xHA sec6-4 swe1Δ bfa1Δ* cells grown overnight in YPD to log phase were released from a G1 phase arrest into fresh media at 25°C. 20 minutes after release the cultures were shifted to 34°C to inactivate sec6-4 and samples were collected at the indicated time points to assay levels of Clb2.

To further test the model, we analyzed the effects of deleting the *BFA1* gene, which encodes a GAP thought to inhibit the MEN via inhibition of Tem1 (Pereira *et al*., 2000; Wang *et al*., 2000; Ro *et al*., 2002). Loss of *BFA1* partially abrogated the late mitotic arrest in *sec6-4 swe1Δ* cells (**Figure 6B**).

These results suggest that the late mitotic arrest caused by *sec6-4* is due to a failure to activate the MEN, consistent with a model in which the events of bud growth generate signals that promote Net1 phosphorylation and are required for full activation of the MEN.

### Components of the TORC2 signaling network are required for hyperphosphorylation of Net1

The discovery that inactivation of Sec6 causes a failure in hyperphosphorylation of Net1, as well as a late mitotic arrest that is dependent upon binding of Net1 to Cdc14, is consistent with a model in which activation of the MEN requires growth-dependent signals that report on the extent of bud growth. However, the data do not rule out alternative models. A key question concerns the nature of the signals that drive Net1 hyperphosphorylation. However, these signals are poorly understood. Over 90 phosphorylation sites have been detected on Net1 and multiple kinases have been implicated (Shou and Deshaies, 2002; Azzam *et al*., 2004; Mah *et al*., 2005; Stark *et al*., 2010; Zhou *et al*., 2021). Mitotic Cdk1 activity is thought to contribute to Net1 phosphorylation; however, only 10 of the many sites that have been detected on Net1 correspond to the minimal Cdk1 consensus site (S/TP). Furthermore, a previous study showed that mitotic spindle assembly, chromosome segregation, and mitotic spindle elongation occur normally in *sec6-4 swe1Δ* cells, despite the failure in bud growth (Anastasia *et al*., 2012). Since all these key events of mitosis require mitotic Cdk1 activity, the fact that they occur normally in *sec6-4 swe1Δ* cells that show a complete failure in Net1 hyperphosphorylation suggests that hyperphosphorylation of Net1 is not a consequence of mitotic Cdk1 activity. Another kinase that has been proposed to phosphorylate Net1 is the Cdc5/Polo kinase; however, mutation of sites thought to be phosphorylated by Cdc5 has no effect on release of Cdc14 or mitotic progression in vivo (Shou and Deshaies, 2002; Zhou *et al*., 2021). Additionally, Net1 hyperphosphorylation induced by Cdc5 overexpression is dependent upon Tem1, which suggests that Cdc5 could influence Net1 phosphorylation via Tem1-dependent activation of other kinases (Visintin *et al*., 2003). No kinase has been shown to be required for the extensive hyperphosphorylation of Net1 seen in vivo. Although kinases have been identified that are capable of disrupting the Net1-Cdc14 complex in vitro it remains unclear whether they work directly to disrupt the complex in vivo (Shou and Deshaies, 2002; Azzam *et al*., 2004).

To investigate further we first tested whether kinases previously proposed to phosphorylate Net1 are required for Net1 phosphorylation in vivo. Cells harboring an analog-sensitive allele of *CDC5* (*cdc5-as1*) were released from a G1 phase arrest and analog inhibitor was added 30 minutes after release. Inhibition of cdc5-as1 caused cells to arrest with hyperphosphorylated Net1^Δ328^, which indicates that Cdc5 is not required for Net1 hyperphosphorylation (**Figure S1A**). Similarly, inhibition an analog-sensitive allele of the *CDC15* kinase (*cdc15-as1*), a key component of the MEN signaling cascade, had no effect on Net1^Δ328^ phosphorylation in cells released from a G1 phase arrest into medium containing benomyl (**Figure S1B)**. Since Cdc15 is required for activation of Dbf2, these results indicate that none of the key kinases in the MEN are required for hyperphosphorylation of Net1, which suggests that the hyperphosphorylation of Net1 that can be detected via electrophoretic mobility shifts is not due to MEN activity. This is consistent with previous studies that found that the MEN is not active in metaphase-arrested cells (Fesquet *et al*., 1999; Jaspersen and Morgan, 2000; Hu *et al*., 2001; Visintin and Amon, 2001; Hu and Elledge, 2002). The results do not rule out the possibility that MEN kinases phosphorylate a small number of sites that do not influence the electrophoretic mobility of Net1.

The evidence for links between Net1 phosphorylation and bud growth suggest that signals associated with cell growth could play a role in hyperphosphorylation of Net1. We therefore tested whether signals known to be associated with cell growth are required for hyperphosphorylation of Net1. Previous work has shown that TOR kinases play important roles in control of cell growth. The TOR kinases are assembled into two distinct multiprotein complexes, referred to as Target of Rapamycin Complexes 1 and 2 (TORC1 and TORC2) (Loewith and Hall, 2011). TORC1 is best understood because it can be inhibited with rapamycin and plays roles in control of ribosome biogenesis, amino acid biosynthesis, and autophagy. A key target of TORC1 is the conserved Sch9 kinase, which controls transcription of ribosome biogenesis genes. To test whether TORC1 is required for hyperphosphorylation of Net1 we synchronized cells and added rapamycin shortly after bud emergence. Rapamycin had no effect on hyperphosphorylation of Net1^Δ328^ in cells released from a G1 phase arrest into medium containing benomyl (**Figure S1C**). Similarly, inhibition of an analog-sensitive version of Sch9 (*sch9-as*) had no effect on hyperphosphorylation of Net1^Δ328^ (**Figure S1D**).

Recent studies have shown that TORC2 activity is correlated with growth rate and the extent of growth. Moreover, a signaling network that surrounds TORC2 is required for normal control of cell size and for nutrient modulation of cell size (Alcaide-Gavilán *et al*., 2018; Lucena *et al*., 2018; Leitao *et al*., 2019). A key downstream target of TORC2 that is required for normal control of cell growth and size is a pair of redundant kinase paralogs called Ypk1 and Ypk2, which are the budding yeast homologs of mammalian SGK kinases (Niles *et al*., 2012). Inhibition of an analog-sensitive allele of *YPK1* in a *ypk2Δ* background (*ypk1-as ypk2Δ*) caused a substantial failure in hyperphosphorylation of Net1^Δ328^ in cells released from a G1 phase arrest into medium containing benomyl (**Figure 7)**. A previous study found that inhibition of Ypk1/2 does not block bud growth (Clarke *et al*., 2017). Together, these observations suggest that the TORC2 signaling network could play a role in the signals that drive hyperphosphorylation of Net1, consistent with the mass spectrometry results showing that multiple components of the TORC2 signaling network are rapidly regulated in response to a shift from rich to poor carbon during mitosis (Table 1).

**Figure 7.**
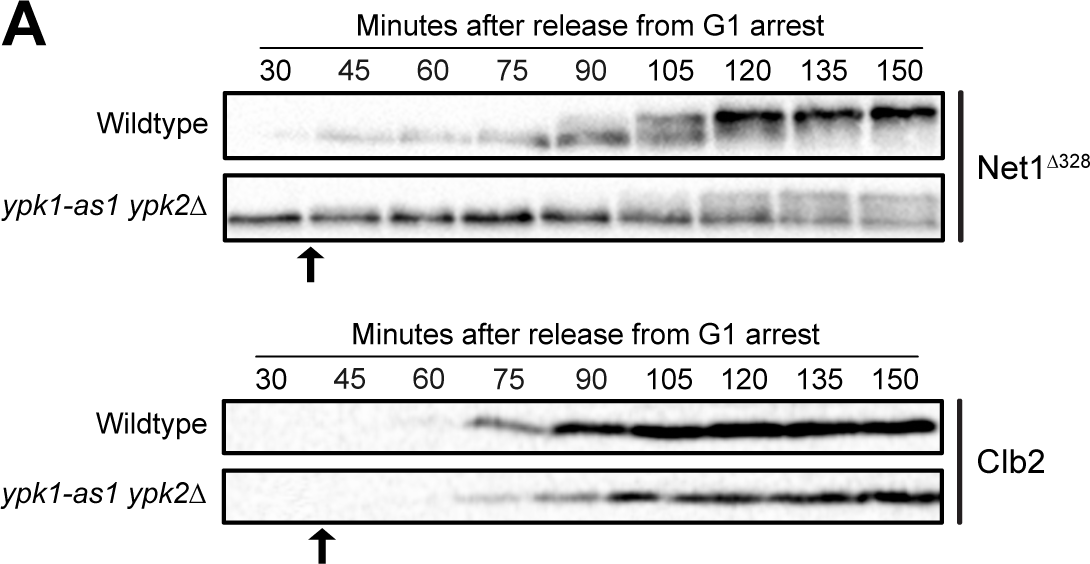
– Ypk1/2 is required for normal hyperphosphorylation of Net1 *in vivo*. *NET1^Δ328^-3xHA* and *NET1^Δ328^-3xHA ypk2Δ ypk1-as1* cells grown overnight in YPD were released from a G1 phase arrest at 25°C. 3MOB-PP1 was added to both cultures to a final concentration of 10 uM at 40 minutes after release. Samples were taken at the indicated time intervals to assay for Net1^Δ328^-3xHA and Clb2 by western blot.

## Discussion

### Numerous components of the MEN are controlled by nutrient-dependent signals

In budding yeast, most cell growth occurs during mitosis as the daughter bud undergoes rapid expansion (Leitao and Kellogg, 2017). Furthermore, bud growth is required for progression through mitosis (Anastasia *et al*., 2012; Jasani *et al*., 2020). Previous studies led us to hypothesize that the extent of bud growth is measured to ensure that an appropriate amount of growth occurs, and that the threshold amount of growth required for mitotic progression is reduced in poor nutrients, which would explain why daughter cells are born at a very small size in poor nutrients (Anastasia *et al*., 2012; Jasani *et al*., 2020). Testing this hypothesis will require identification of molecular mechanisms that link mitotic progression to bud growth, as well as mechanisms that set the threshold amount of bud growth required for cell cycle progression. Here, we searched for molecular mechanisms that link completion of mitosis to bud growth. To do this, we took advantage of the fact that changes in nutrient availability rapidly re-set the threshold amount of growth required for cell cycle progression (Fantes and Nurse, 1977; Leitao and Kellogg, 2017). We reasoned that proteins involved in measuring cell growth and setting the growth threshold should show rapid changes in phosphorylation in response to a shift to poor nutrients. Thus, we used proteome-wide mass spectrometry to identify proteins that undergo rapid changes in phosphorylation when cells in mitosis are shifted from rich to poor carbon. This revealed that numerous components of the MEN are regulated by nutrient availability. Since activation of MEN signaling initiates exit from mitosis, the MEN is well-positioned to receive physiological signals related to nutrients and cell growth to ensure that mitotic exit is initiated only when an appropriate amount of growth has occurred. A role for the MEN in controlling mitotic exit in response to nutrient and growth cues would be compatible with previously proposed models in which migration of the daughter nucleus into the daughter bud promotes mitotic exit. For example, signals associated with cell growth and nutrient availability could influence initiation of mitotic exit, while migration of the nucleus into the daughter bud could initiate positive feedback that fully activates the MEN to complete mitotic exit.

An alternative hypothesis could be that a shift to poor carbon reduces the availability of ATP, leading to a general slowdown in cellular events and a decrease in the rate of phosphorylation of proteins. Several observations argue against this interpretation. First, a shift to poor carbon late in mitosis does not cause a delay in mitotic exit, which suggests that the shift does not cause a non-specific slowdown of mitotic events (Leitao and Kellogg, 2017). Rather, a shift to poor carbon in late mitosis may not cause a delay because sufficient bud growth has already occurred, especially as the shift should immediately lower the threshold amount of growth required for cell cycle progression (Fantes and Nurse, 1977). A further argument against a general reduction in ATP is that the top proteins that underwent dephosphorylation are enriched in proteins involved in the MEN, TORC2 signaling, ribosome biogenesis and membrane trafficking, all of which are involved in growth or cell cycle progression and would be expected to change when growth rate is slowed by poor nutrients. In contrast, few proteins involved in mitotic spindle dynamics, which are known to undergo phosphorylation during mitosis, were identified.

### Hyperphosphorylation of Net1 occurs before mitotic exit

Among the MEN components identified by mass spectrometry, Net1 was amongst the most prominent with 11 phosphorylation sites that showed substantial loss of phosphorylation. Net1 prevents mitotic exit by binding and inhibiting the Cdc14 phosphatase in the nucleolus. The dissociation of Cdc14 from Net1 initiates mitotic exit as well as a positive feedback loop that promotes mitotic exit. It is thought that hyperphosphorylation of Net1 promotes dissociation of Cdc14, thereby initiating mitotic exit, although there is evidence that hyperphosphorylation of Net1 may not be sufficient to release Cdc14 (Traverso *et al*., 2001; Visintin *et al*., 2003; Azzam *et al*., 2004; Zhou *et al*., 2021). A previous study found that Cdc14 is not released from the nucleus in metaphase-arrested cells, consistent with the idea that phosphorylation of Net1 is not sufficient to drive release of Cdc14 from the nucleus (Stegmeier *et al*., 2002). Here, we found that Net1 undergoes full hyperphosphorylation in metaphase-arrested cells, which indicates that Net1 hyperphosphorylation is not strictly correlated with mitotic exit and does not depend upon the events of mitotic exit. The finding that Net1 is fully hyperphosphorylated in metaphase arrested cells is consistent with previously proposed models in which hyperphosphorylation of Net1 is necessary but not sufficient to initiate exit from mitosis.

### Phosphorylation of Net1 is correlated with daughter bud growth

What are the physiological signals that drive hyperphosphorylation of Net1 before mitotic exit? Several observations suggest that signals associated with cell growth influence Net1 phosphorylation. First, hyperphosphorylation of Net1 occurs gradually during bud growth, which indicates that Net1 phosphorylation is correlated with bud growth. Furthermore, we found that metaphase-arrested cells undergo unusually extensive bud growth, which leads to abnormally large daughter buds as well as unusually extensive hyperphosphorylation of Net1. Similarly, we used growth in poor carbon sources to show that reducing the rate and extent of bud growth leads to a corresponding reduction in the rate at which hyperphosphorylated forms of Net1 accumulate as well as a reduction in the maximal extent of Net1 phosphorylation. Thus, hyperphosphorylation of Net1 is strongly correlated with the rate and extent of bud growth, which suggests that the events of growth generate signals that help drive hyperphosphorylation of Net1.

### Hyperphosphorylation of Net1 is dependent upon membrane trafficking events that drive bud growth

Further evidence for a link between Net1 phosphorylation and cell growth came from our finding that hyperphosphorylation of Net1 is dependent upon membrane trafficking events that drive bud growth. In *sec6-4 swe1Δ* cells the initial steps of mitotic exit occur normally as the spindle elongates during anaphase (Anastasia *et al*., 2012), yet hyperphosphorylation of Net1 fails to occur. These observations provide further evidence that hyperphosphorylation of Net1 is closely associated with bud growth, rather than with early steps of anaphase and mitotic exit. Previous studies have shown that an arrest of membrane trafficking events that are required for plasma membrane growth leads to inhibition of ribosome biogenesis (Mizuta and Warner, 1994; Li *et al*., 2000). Therefore, the failure in Net1 phosphorylation in *sec6-4 swe1Δ* cells could be a more direct consequence of an arrest of ribosome biogenesis, although it is unclear from previous studies whether arresting membrane growth causes an arrest of ribosome biogenesis on a sufficiently rapid time scale to account for the observed effects on Net1 phosphorylation. It is also possible that the inhibition of ribosome biogenesis caused by blocking membrane trafficking events is due to a failure in Net1 phosphorylation, as previous studies have shown that Net1 is required for normal transcription of rRNA genes (Shou *et al*., 2001; Hannig *et al*., 2019).

The fact that Net1 is intimately connected to ribosome biogenesis, a major facet of cell growth, further suggests a potential connection between Net1 hyper-phosphorylation and cell growth (Straight *et al*., 1999; Shou *et al*., 2001; Hannig *et al*., 2019). An interesting possibility is that gradual phosphorylation of Net1 during bud growth ensures that the rate of rDNA gene transcription increases during growth to meet the increased protein synthesis requirements of larger cells. Consistent with this, a previous study found that phosphorylation of Net1 promotes binding and activation of RNA polymerase PolI (Hannig *et al*., 2019). This would suggest a model in which the events of growth generate feedback signals that promote further growth and also provide a measure of the extent of growth.

### Bypassing the MEN relieves a late mitotic arrest caused by a failure in bud growth

In previous work, we found that *sec6-4 swe1Δ* cells arrest in late anaphase with long mitotic spindles, separated nuclei, and high levels of mitotic cyclin, which indicates that they fail to complete mitotic exit (Anastasia *et al*., 2012). The cause of this late mitotic arrest was unknown. Here, we found that the late anaphase arrest in *sec6-4 swe1Δ* cells can be bypassed by preventing Net1-dependent inhibition of Cdc14. Together, these observations suggest that the late mitotic arrest caused by blocking bud growth is due to a failure to activate the MEN, which is consistent with the hypothesis that activation of the MEN is dependent upon bud growth.

### Net1 undergoes extensive hyperphosphorylation that is not dependent upon key components of the MEN

Net1 is one of the most highly phosphorylated proteins in budding yeast. Various mass spectrometry analyses have identified 91 phosphosites. Of the 11 sites identified in our analysis, 10 were identified in previous analyses, while 1 was not previously identified.

The functions of Net1 phosphorylation remain poorly understood. Many of the 91 sites were identified in non-quantitative proteome-wide analyses, which makes it unclear when they are phosphorylated in a cell cycle-dependent manner or whether they are phosphorylated at significant stoichiometries that are likely to be biologically relevant. A Net1 mutant in which all 91 sites were mutated (*net1-91A*) is viable and causes only a slight delay in mitotic exit (Zhou *et al*., 2021). However, whether the *net1-91A* mutant eliminates phosphorylation-induced electrophoretic mobility shifts in vivo was not tested, so it is unclear whether relevant sites were mutated. Furthermore, mutating 91 phosphorylation sites almost certainly eliminates regulation by multiple kinases, which makes interpretation of the results difficult. For example, if Net1 plays both positive and negative roles in controlling ribosome biogenesis and mitotic exit, as suggested by previous studies, elimination of 91 phosphorylation sites could cause both gain- and loss-of-function effects that cancel each other out (Straight *et al*., 1999; Shou *et al*., 2001; Hannig *et al*., 2019). The fact that mutating subsets of sites appears to cause stronger effects is consistent with this possibility (Zhou *et al*., 2021).

Previous studies suggested that Cdc5 and Dbf2, two key components of the MEN, could phosphorylate Net1 (Shou and Deshaies, 2002; Yoshida and Toh-e, 2002; Visintin *et al*., 2003; Mah *et al*., 2005; Zhou *et al*., 2021). Previous studies mapped phosphorylation sites on Net1 that are dependent upon Cdc5 or Cdc15. In some cases, mutation of these sites causes mitotic delays, especially in sensitized genetic backgrounds (Loughrey Chen *et al*., 2002; Shou and Deshaies, 2002; Shou *et al*., 2002; Zhou *et al*., 2021). However, the phosphorylation site mutants do not cause a failure in mitotic exit or a loss of viability, and they appear to have only mild effects in an otherwise wildtype background, which indicates that they are not essential for mitotic exit. No previous studies have tested whether Cdc5, Cdc15 or Dbf2 are required for hyperphosphorylation of Net1 that can be detected via electrophoretic mobility shifts.

Here, we found that Cdc5 and Cdc15 are not required for hyperphosphorylation of Net1 in vivo. Since Cdc15 is required for activation of Dbf2, these findings suggest that none of the protein kinases in the MEN are required for the extensive hyperphosphorylation of Net1 that can be detected via electrophoretic mobility shifts. Previous studies found that key components of the MEN are inactive during a metaphase arrest, which provides further evidence that hyperphosphorylation of Net1 in metaphase-arrested cells is not due to signaling from components of the MEN (Fesquet *et al*., 1999; Jaspersen and Morgan, 2000; Hu *et al*., 2001; Visintin and Amon, 2001; Hu and Elledge, 2002). Furthermore, the fact that full hyperphosphorylation of Net1 occurs even when mitotic spindle assembly is blocked shows that there is extensive hyperphosphorylation of Net1 that is not dependent upon any events associated with the function or orientation of the mitotic spindle. A potential explanation for these observations is that hyperphosphorylation of Net1 is the consequence of early signals that are necessary but not sufficient for initiating full activation of the MEN signaling network.

### TORC2 signaling is required for growth-dependent hyperphosphorylation of Net1

Since hyperphosphorylation of Net1 appears to be correlated with growth and dependent upon growth, we tested whether it is influenced by the activity of TORC1 or TORC2, which play essential roles in growth control. Inactivation of TORC1 with rapamycin had no effect on Net1 hyperphosphorylation. Similarly, inactivation of Sch9, a key target of TORC1 that controls ribosome biogenesis, also had no effect on the extent of Net1 phosphorylation. In contrast, inactivation of Ypk1 and Ypk2, key targets of TORC2 that are homologs of mammalian SGK kinases, caused a nearly complete failure in Net1 hyperphosphorylation.

The discovery that components of the TORC2 network are required for Net1 phosphorylation is particularly interesting. Multiple components of the TORC2 network were identified as top hits in our mass spectrometry analysis of the effects of a shift from rich to poor carbon during mitosis. These include core components of the TORC2 complex (Avo2, Avo1 and Tsc11) as well as a key downstream effector, Ypk1. Furthermore, previous studies have shown that Ypk1/2 and their surrounding signaling network are required for normal control of cell size and for nutrient modulation of cell size (Lucena *et al*., 2018). For example, decreased activity of Ypk1/2 causes a large decrease in cell size. Ypk1/2 activate ceramide synthase, which produces signaling lipids required for normal control of cell size. An inhibitor of the ceramide synthesis pathway causes a dose-dependent decrease in cell size, and loss of ceramide synthase causes a complete failure of nutrient modulation of TORC2 signaling and cell size. TORC2 signaling, as measured via phosphorylation of Ypk1/2, is modulated by nutrient availability and increases throughout the interval of bud growth (Alcaide-Gavilán *et al*., 2018; Lucena *et al*., 2018). Finally, loss of Rts1, a conserved PP2A regulatory subunit that influences TORC2 signaling, causes a complete breakdown in the relationship between cell size and growth rate during bud growth in mitosis (Leitao *et al*., 2019). Together, these observations suggest that gradually rising TORC2 signaling during bud growth could drive growth-dependent phosphorylation of Net1, and that nutrient modulation of TORC2 signaling could explain the effects of nutrient availability on Net1 phosphorylation and cell size at completion of mitosis.

#### What are the signals that drive mitotic exit?

A puzzling aspect of the MEN has been that Cdc14, which plays perhaps the most critical role in initiation of mitotic exit, is conserved across eukaryotic cells, yet the role of Cdc14 in promoting mitotic exit is not conserved, as budding yeast is the only known organism in which Cdc14 functions in mitotic exit. For example, the fission yeast homolog of Cdc14 appears to function in a pathway that controls mitotic entry (Trautmann *et al*., 2001). A potential explanation could come from the fact that budding yeast is the only known organism in which most cell growth occurs during mitosis. Thus, it seems possible that Cdc14 has been co-opted in budding yeast to function in an unusual pathway that links mitotic exit to bud growth. Our results support this model, as they suggest that the MEN responds to growth-dependent signals that define the duration and extent of bud growth in late mitosis. The data do not rule out previously suggested models in which the MEN ensures that mitotic exit does not occur until the daughter nucleus has entered the bud. Indeed, the existence of a mechanism to ensure that nuclear segregation occurs only when the bud has become sufficiently large to accommodate the daughter nucleus would appear to be essential for yeast cells that rely on a highly unusual growth mode (i.e. bud growth in mitosis) to reproduce.

## Materials and Methods

### Strain construction, media, and reagents

All strains are in the W303 background (MATa leu2-3,112 ura3-1 can1-100 ade2-1 his3-11,15 trp1-1 GAL+ ssd1-d2). Additional genetic features are listed in Table 2. Gene deletions and epitope tagging were performed by standard PCR amplification and homologous recombination (Longtine et al. 1998; Janke et al. 2004; Lee et al. 2013).

**Table 2:**
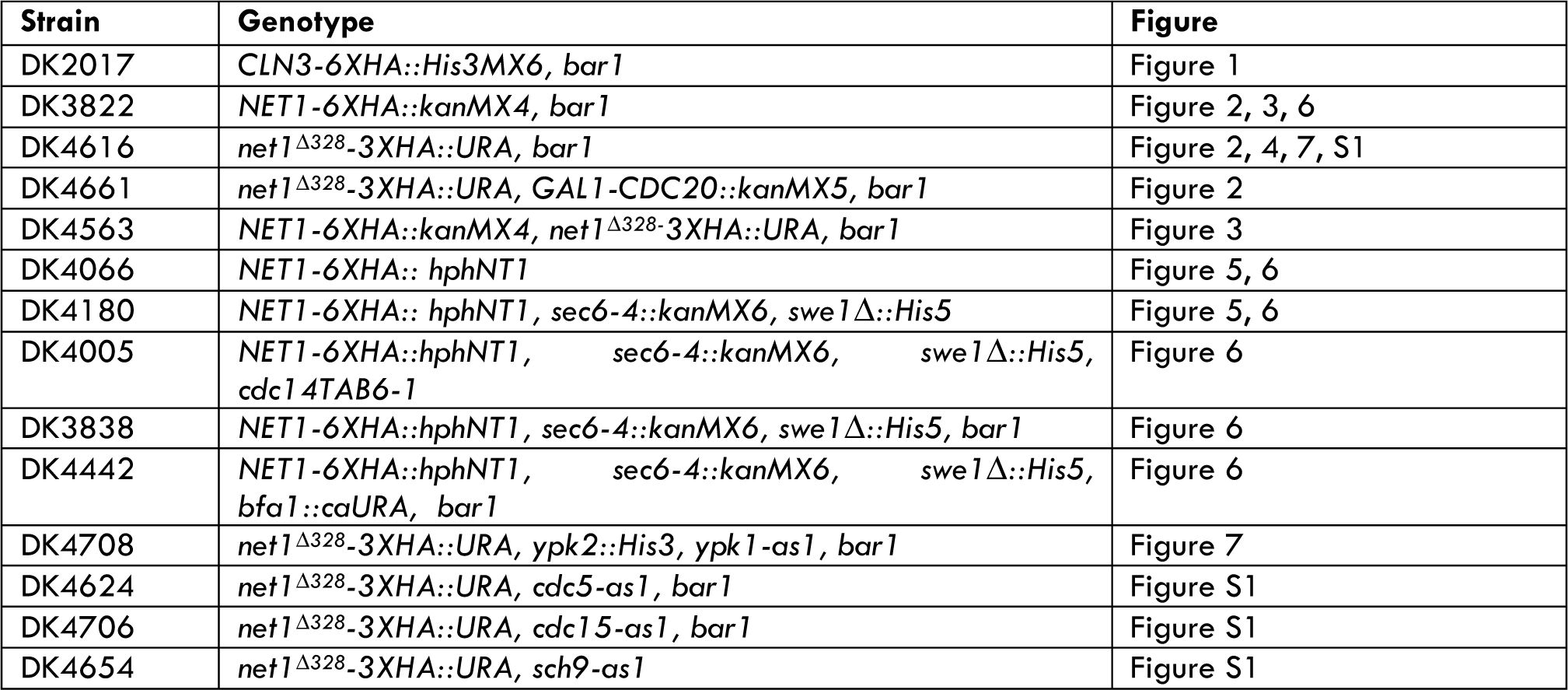
Strains used in this study.

| Strain | Genotype | Figure |
| --- | --- | --- |
| DK2017 | CLN3-6XHA::His3MX6, bar1 | Figure 1 |
| DK3822 | NET1-6XHA::kanMX4, bar1 | Figure 2, 3, 6 |
| DK4616 | net1<sup>Δ328</sup>-3XHA::URA, bar1 | Figure 2, 4, 7, S1 |
| DK4661 | net1<sup>Δ328</sup>-3XHA::URA, GAL1-CDC20::kanMX5, bar1 | Figure 2 |
| DK4563 | NET1-6XHA::kanMX4, net1<sup>Δ328</sup>-3XHA::URA, bar1 | Figure 3 |
| DK4066 | NET1-6XHA::hphNT1 | Figure 5, 6 |
| DK4180 | NET1-6XHA::hphNT1, sec6-4::kanMX6, swe1Δ::His5 | Figure 5, 6 |
| DK4005 | NET1-6XHA::hphNT1, sec6-4::kanMX6, swe1Δ::His5, cdc14TAB6-1 | Figure 6 |
| DK3838 | NET1-6XHA::hphNT1, sec6-4::kanMX6, swe1Δ::His5, bar1 | Figure 6 |
| DK4442 | NET1-6XHA::hphNT1, sec6-4::kanMX6, swe1Δ::His5, bfa1::caURA, bar1 | Figure 6 |
| DK4708 | net1<sup>Δ328</sup>-3XHA::URA, ypk2::His3, ypk1-as1, bar1 | Figure 7 |
| DK4624 | net1<sup>Δ328</sup>-3XHA::URA, cdc5-as1, bar1 | Figure S1 |
| DK4706 | net1<sup>Δ328</sup>-3XHA::URA, cdc15-as1, bar1 | Figure S1 |
| DK4654 | net1<sup>Δ328</sup>-3XHA::URA, sch9-as1 | Figure S1 |

Strains were grown in YEP medium (1% yeast extract, 2% peptone, and 40 mg/mL adenine) containing 2% dextrose (YPD), 2% galactose (YPGal), or 2% glycerol/2% ethanol (YPG/E), as indicated in the text. In experiments involving analog-sensitive alleles of kinases, cells were grown in media that was not supplemented with adenine.

The Net1^Δ328^ phosphorylation reporter was created by amplifying a region of the *NET1* gene that includes the promoter and the first 806 amino acids (oligos: ATACC-CGGGCGTAGGGAGCGATATGTGCATTATG and ATAGGTACCCTTCTTGTTTGAA-TTCGGATAAAAGCTTTTCGCC). The resulting PCR fragment was cloned into the KpnI and XmaI sites of pDK51, which expresses proteins with a C-terminal 3XHA tag and carries the *URA3* marker, to create pDK131. The plasmid is cut with StuI to target integration at the *URA3* gene.

Benomyl was solubilized in 100% DMSO to prepare a 20 mg/mL stock solution. Media containing benomyl was prepared by first heating the media to 100°C before adding the drug to a final concentration of 30 ug/mL. The media was then allowed to cool at room temperature while stirring.

### Cell cycle time courses and western blotting

For cell cycle time courses, cultures were grown overnight at room temperature to log phase (OD_600_ = 0.4 - 0.7). The cells were then arrested in G1 phase by the addition of 0.5 ug/mL or 15 ug/mL alpha factor for *bar1* or BAR1 strains, respectively. Cultures were arrested for 3 to 3.5 hours and then released from the arrest by three washes in fresh media not containing mating pheromone. All time courses were performed at 25°C, except for experiments involving *sec6-4* strains, which were carried out at 34°C. Alpha factor was added back to cultures 60 minutes after release to prevent initiation of a second cell cycle. At each time point, 1.6 mL of sample were collected in screw cap tubes and centrifuged at 15,000 rpm for 15 seconds. The supernatant was then removed and 200 uL of acid-washed glass beads were added before freezing samples in liquid nitrogen. Cell pellets were lysed in 140 uL 1x SDS-PAGE sample buffer (65 mM Tris-HCl pH 6.8, 3% SDS, 10% glycerol, 100 mM {²-glycerophosphate, 50 mM NaF, 5% {²-mercaptoethanol, 3 mM PMSF, and bromophenol blue) by bead beating in a Biospec Mini-Beadbeater-16 at 4°C for 2 minutes. Lysed samples were spun down in a microcentrifuge for 15 seconds at 15,000 rpm before incubating in a 100°C water bath for 6 minutes followed by centrifugation at 15,000 rpm for 10 minutes.

SDS-PAGE was carried out as previously described (Harvey *et al*., 2011). 10% polyacrylamide gels with 0.13% bis-acrylamide were used for analysis of Net1, Clb2, and Nap1 (loading control). Blots were probed with the primary antibody at 1–2 ug/ml at 4°C overnight in PBST (phosphate-buffered saline, 250 mM NaCl, and 0.1% Tween-20) containing 4% nonfat dry milk. Primary antibodies used to detect Clb2 and Nap1 were rabbit polyclonal antibodies generated as described previously (Kellogg and Murray, 1995; Sreenivasan and Kellogg, 1999; Mortensen *et al*., 2002). Net1-6xHA and Net1^Δ328^-3xHA were detected by a mouse monoclonal antibody (12CA5). Primary antibodies were detected by an HRP-conjugated donkey anti-rabbit secondary antibody (#NA934V; GE Healthcare) or HRP-conjugated donkey anti-mouse secondary antibody (#NXA931; GE Healthcare) incubated in PBST containing 4% nonfat dry milk for 1 hr at room temperature. Blots were rinsed in PBS before detection via chemiluminescence using ECL reagents (#K-12045-D50; Advansta) with a Bio-Rad (Hercules, CA) ChemiDoc imaging system.

### Cell size analysis

For Figure 4B, cell cultures were grown overnight at room temperature in YPD, YPGal, or YPG/E to an OD_600_ between 0.4 and 0.6. During the time course, 450 ul of culture was collected at each time point and fixed by addition of 50 uL of 37% formaldehyde to the culture medium followed by incubation at room temperature for 1 hour. Cells were then pelleted and resuspended in 0.25 ml PBS containing 0.02% sodium azide and 0.1% Tween-20. Cell size was measured on the same day using a Coulter Counter (Coulter Counter Z2; Beckman, Fullerton, CA).

### Tandem mass spectrometry analysis

Cells were grown overnight to log phase in YPD medium and were then arrested in G1 phase with alpha factor. The cells were released from the arrest into fresh YPD medium and at 90 minutes after release one half of the culture was shifted to poor carbon (YPG/E) by pelleting the cells and resuspending in YPG/E three times. At ten minutes after the first wash into poor carbon, the cells from 50 mls of both cultures (O.D. 600 of 0.5) were pelleted in a 50 ml conical tube, resuspended in 1 ml of media, transferred to a new 1.6 ml screw top, and then pelleted again for 15 sec in a microfuge. After removing the supernatant, 200 uL of acid-washed beads were added and the samples were frozen on liquid nitrogen. Parallel samples were taken for western blotting to ensure that the shift to poor carbon occurred at peak Clb2 levels. A total of 2 biological replicates were carried out and analyzed by mass spectrometry. A strain carrying Cln3-6XHA (DK2017) was used for these experiments because we wanted to test whether the Cln3 protein present in mitosis responds rapidly to a shift to poor carbon. Western blotting showed that Cln3 rapidly disappeared when the cells were shifted to poor carbon within 5 minutes, as observed previously for Cln3 in G1 phase.

Cell pellets were lysed in a Biospec Mini-Beadbeater-16 at 4°C in 500 uL of Lysis Buffer (8M urea, 75 mM NaCl, 50 mM Tris, pH 8.0, 50 mM B-glycerolphosphate, 1 mM NaVO_3_, 10 mM sodium pyrophosphate, 1 mM PMSF) by two 1-minute rounds of bead-beating at 4°C, with 1 minute of chilling in an ice water bath in between. The lysate was then centrifuged at 14,000 rpm at 4°C for 10 minutes and the supernatant was transferred to a new tube and frozen on liquid nitrogen. The samples were prepared for quantitative proteome and phosphoproteome profiling following the SL-TMT protocol that includes the “mini-phos” phosphopeptide enrichment strategy (Navarrete-Perea, Yu et al. 2018).

## Acknowledgements

We thank the Shokat lab for providing 3MOB-PP1. We also thank the members of the Kellogg lab for advice and critical reading of the manuscript. This work was supported by NIH grant R35 GM131826.

## Supplementary Figure Legends

**Figure S1.**
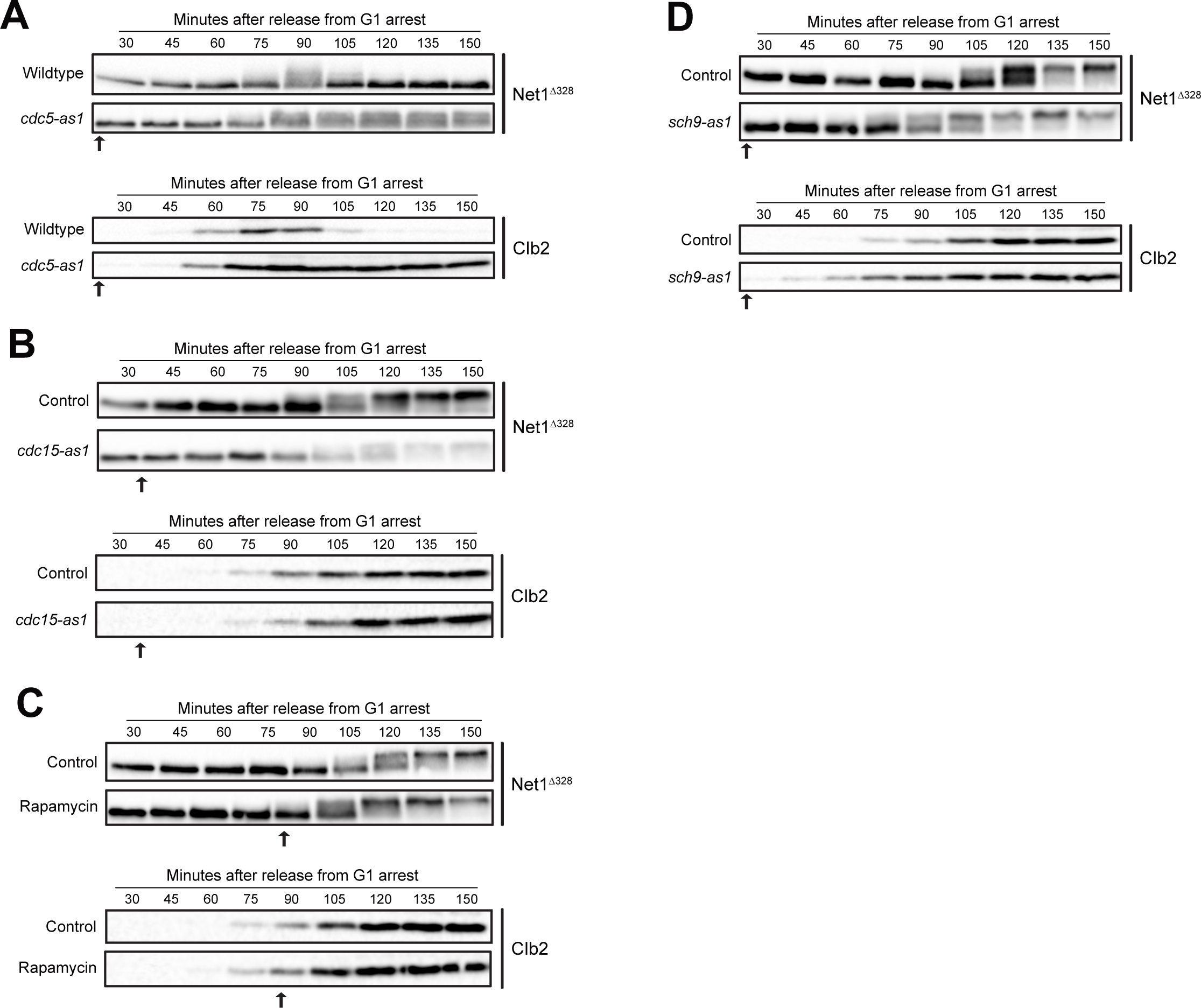
– Analysis of candidate Net1 kinases *in vivo.* For all experiments, cells were released from a G1 phase arrest and inhibitor was added at the times indicated by the arrow. Samples were then taken at the indicated time points to assay for *NET1^Δ328^* and Clb2 by western blot. (A) *NET1^Δ328^-3xHA* and *net1^Δ328^-3xHA cdc5-as1* cells were released from G1 phase arrest into fresh media at 30°C. 10 uM CMK was added 30 minutes after release. (B) *NET1^Δ328^-3xHA* and *NET1^Δ328^-3xHA cdc15-as1* cells were released from G1 phase arrest into fresh media containing benomyl at 25°C. 10 uM 1-NA-PP1 was added 40 minutes after release. (C) *NET1^Δ328^-3xHA* cells were released from G1 phase arrest into fresh media containing benomyl 25°C. The culture was split into two aliquots and 0.22 uM rapamycin was added to one aliquot 90 minutes after release. (D) *NET1^Δ328^-3xHA* and *NET1^Δ328^-3xHA sch9-as1* cells released from G1 phase into fresh media containing benomyl at 25°C. 5 uM 1-NM-PP1 was added 30 minutes after release.

## Notes

### Competing Interest Statement

The authors have declared no competing interest.

